# SHYBRID: A graphical tool for generating hybrid ground-truth spiking data for evaluating spike sorting performance

**DOI:** 10.1101/734061

**Authors:** Jasper Wouters, Fabian Kloosterman, Alexander Bertrand

**Author notes:** This work was carried out at the ESAT Laboratory of KU Leuven, in the frame of KU Leuven Special Research Fund projects C14/16/057, and the Research Foundation Flanders (FWO) project FWO G0D7516N. This project has received funding from the European Research Council (ERC) under the European Unions Horizon 2020 research and innovation programme (grant agreement No 802895). This research received funding from the Flemish Government under the Onderzoeksprogramma Artificile Intelligentie (AI) Vlaanderen programme. The scientific responsibility is assumed by its authors.

## Abstract

Spike sorting is the process of retrieving the spike times of individual neurons that are present in an extracellular neural recording. Over the last decades, many spike sorting algorithms have been published. In an effort to guide a user towards a specific spike sorting algorithm, given a specific recording setting (i.e., brain region and recording device), we provide an open-source graphical tool for the generation of hybrid ground-truth data in Python. Hybrid ground-truth data is a data-driven modelling paradigm in which spikes from a single unit are moved to a different location on the recording probe, thereby generating a virtual unit of which the spike times are known. The tool enables a user to efficiently generate hybrid ground-truth datasets and make informed decisions between spike sorting algorithms, fine-tune the algorithm parameters towards the used recording setting, or get a deeper understanding of those algorithms.

## I. Introduction

After more than a century of brain research, and despite many breakthroughs, it is still largely unknown how brain activity gives rise to cognition. To aid researchers in getting a better understanding of brain function, many techniques have been developed to look at neuronal activity dynamics. Although recent techniques, such as calcium imaging, are getting increasingly popular, electrophysiology is still indispensable because it has the highest available temporal resolution among other available techniques. Electrophysiological recordings allow researchers to look at brain activity at a time scale that reveals neuronal action potentials in great detail. To increase the spatial extent of such electrophysiological recordings, recently recording hardware has undergone a drastic evolution leading to high-density neural probes [1] [2].

Most commonly, extracellular recordings are performed by inserting an array of electrodes into the brain. Electrodes that are in close proximity to a neuron will pick up a transient potential, referred to as a spike, when the neuron fires an action potential. Such action potentials are believed to be the main mechanism of communication between neurons. By measuring spikes from several neurons, one can try to get a better understanding of information processing in neuronal circuits. However, typically many neurons surround a single electrode, leading to spikes from several neurons to be picked up by the same electrode. On the other hand, in high-density recordings a spike from a single neuron is picked up by multiple electrodes. To resolve the mixture of spikes from several neurons and extract the spike times of the individual neurons embedded in the recordings, a spike sorting [3] [4] algorithm can be applied to the electrophysiological recording. A reliable recovery of the spike trains of individual neurons, enables a wide range of analysis [5] that ultimately advance our understanding of the brain [6] [7] [8]. Furthermore, a better understanding of the brain at the level of single neurons can create clinical impact [9] [10].

Over the last two decades a myriad of spike sorting [11] [12] [13] [14] [15] [16] [17] [18] [19] algorithms have been published. The development of these spike sorting algorithms has been driven by both the availability of novel computational methods [20] [21], as well as the ongoing evolution in the channel count and density of recording equipment [1] [2]. This has led to a broad spectrum of spike sorting algorithms, each with their own (dis)advantages, and often with only subtle differences in their underlying mechanisms. As a result, spike sorting users are often left pondering which algorithm they should use to get the spike sorting job done on their specific data.

The question which algorithm and parameters to use in a particular setting, usually boils down to the question which spike sorting package is the most performing in terms of spike sorting accuracy and computational complexity. Unfortunately, the question related to the spike sorting accuracy is a hard one to answer, due to the lack of thorough comparative studies. In the past, initiatives for spike sorting validation have been set up [22] [23], but they have not led to a systematic validation of recent spike sorting algorithms. One of the reasons for the lack of such comparative studies is the absence of an extensive collection of ground-truth data. Two main paradigms for generating ground-truth data are in use: paired recordings [24] [25] [26] and simulation-based ground-truth data [27] [28] [29] [30] [31]. Both paradigms have their limitations as will be discussed next.

In paired recordings a single neuron is isolated (preferably in vivo) using a high signal-to-noise single cell approach, such as intracellular or juxtacellular recordings, that provides an accurate record of the targeted cell’s spike times. Another recording device, e.g., a neural probe, is then lowered into the brain in close vicinity to the isolated neuron. This probe recording is analysed with a spike sorting algorithm, which can be partially validated using the ground-truth spike times obtained from the intracellularly or juxtacellularly recorded cell. The problem with paired recordings is that they involve an expensive, time consuming procedure. The procedure often fails, as it is very difficult to register the activity of the cell on both recording devices. For the paired recording to succeed, a very precise mutual positioning of both recording devices is required. Note that paired recordings have also been performed in vitro in the context of spike sorting validation [18], which could alleviate some of the experimental difficulties when compared to the in-vivo procedure.

It also remains unclear whether an algorithm’s spike sorting accuracy for data with a specific recording setting is indicative for its accuracy using data recorded under a different setting. The recording setting is defined by both the brain region of interest as well as the recording equipment. Differences between brain regions, e.g., the differences in cell type distribution and cell density, will affect the extracellular recording content. Also different experimental protocols can cause changes in the local spiking activity. Differences in recording equipment, such as different probe channel layouts and channel densities, will naturally lead to a different spike feature space, i.e., the information that is used to assign detected spikes to singleunit spike clusters. The data acquisition system that is used will add electronics noise to the recording. The characteristics of such electronics noise can vary between different models of acquisition systems, or even between different acquisition systems of the same model. All of the above factors will influence the final recording, and as such will also affect the spike sorting results and performance. The recording setting intrinsics thus prevent us from generalizing spike sorting accuracy obtained on paired recordings.

As opposed to paired recordings, ground-truth data generated from computational simulations are more flexible in coping with changing recording settings. The downside of simulated data is that they often lack the richness of real recordings. To ameliorate this problem, background noise can be extracted from real recordings [32], while spikes that are generated from computational neuronal models are superimposed. Although such an approach improves the richness of the data, obtaining realistic simulations that are representative for a specific recording setting requires modelling expert knowledge. This is especially the case if realistic spike timing and scaling statistics are of interest. If realistic variations among spikes from different neurons are desired, morphologically detailed models have to be acquired, a process which is nontrivial and time-consuming. However, several large efforts have been undertaken to provide the community with a public collection of biophysically detailed neuron models [33] [34] [35].

Here, we present an open-source graphical tool which allows spike sorting users to transform their *own* recordings into ground-truth data in an efficient and controlled fashion. The ground truth generation paradigm which is used in our tool, is the so-called hybrid data paradigm that has been used for the validation of some spike sorting algorithms [14] [15]. The generation of such ground-truth data is a delicate process, which at least requires visual feedback on the generated ground truth and ideally some user interaction to make the key decisions during the generation process. Hybrid data does not rely on computational models, nor does it suffer from the complexity of obtaining paired recordings. Hybrid data generation can be seen as a data-driven modelling approach: spike templates, spike timings and scalings are directly extracted from the data. As such, the ground-truth data that is generated is automatically tailored to the user’s recording setting. This ground-truth data can then be used to make an informed decision on which spike sorting algorithm is most suitable for a specific recording setting, fine-tune the algorithm parameters towards the used recording setting, or get a deeper understanding of those algorithms.

The tool takes as an input the raw data and initially curated spike sorting results, e.g., obtained from manual cluster cutting. As a first step a template is calculated for every singleunit spike train. Next, the user visually inspects the template fit for a selection of spikes in the corresponding spike train. If the scaled template model is deemed reasonable, the user moves the entire unit, i.e., the combination of the spike train and its corresponding template, to a different spatial location on the probe. The final step of relocating the unit is key for the relocated unit to be considered as a valid ground-truth unit. For the original unit, one can not make this ground-truth assumption. Indeed, in the initial spike sorting results, one can easily classify a spike cluster as originating from a single neuron, but it is a lot harder to show that all spikes from the specific neuron are present in the cluster. Such missed detections (false negatives) in the initial sorting might be erroneously interpreted as false positives when the original recording and initial spike sorting are used for the validation of other (and potentially better) spike sorters. By changing the spatial location of the hybrid unit with respect to the original unit, this problem is bypassed because the spatial footprint of the hybrid unit is different now from the spatial footprint of the original unit, and as such potential false negatives in the initial sorting are very unlikely to interfere with the hybrid unit in terms of spike sorting.

A main advantage of the presented tool is the visual feedback that is provided throughout the hybrid data generation. The visual feedback allows a user to closely monitor and control the hybrid data quality. For example, visual feedback of the spike rate of every individual channel allows a user to control the spike sorting difficulty of the hybrid unit. Inserting a hybrid unit into a silent spatial region of the probe will likely be easier to sort than inserting a hybrid unit in a very active spatial region. The tool is also capable of fully automatic hybrid ground-truth data generation for use cases where less control is acceptable. An efficient method is included to validate spike sorting results obtained on the generated hybrid ground-truth data. The tool is easy to use and freely available^1^, which will enable a broad audience to generate recording setting-specific ground-truth data for improving their spike sorting related research.

In this work we will focus on generating hybrid data from high-density neural probe recordings. However, the tool is by no means limited to neural probes, it can be used without any modification on various other multi-channel recording devices, e.g., microelectrode arrays (MEAs) [36] [37]. This work as a whole might also be useful outside of the field of neuronal spike sorting. The validation of sEMG [38] decomposition algorithms [39] [40], i.e., algorithms for sorting motor unit action potentials that are measured using electrode arrays placed on the skin on top of muscle tissue, might also benefit from the availability of hybrid ground-truth data.

The hybrid ground-truth model is discussed in depth in the following section. In Section III a graphical tool is introduced that implements this hybrid model. Section IV contains a case study where ground-truth data generated with the user interface is used to compare and study spike sorting algorithms. Finally, the proposed method and results are discussed in Section V.

## II. Hybrid ground-truth model

Before introducing the graphical tool for spiking hybrid ground-truth data generation, we will first introduce and formalize our hybrid ground-truth model. The basic idea behind the generation of hybrid data [14] [15] is to remove singleunit spikes from one location of the probe and re-introduce them at another location with an additional temporal offset. We thus create a fictitious neuron of which the spike waveforms are modelled as scaled spike templates estimated from the initial spike sorting results. This unit is then injected at another location on the probe to prevent false negatives (i.e. spikes from the original neuron that were not identified) in the initial spike sorting results from being wrongly interpreted as false positives when later using the hybrid data for evaluating spike sorting performance. This spatial migration is sufficient for decoupling the fictitious neuron from its donor neuron for spike sorting purposes. This approach depends on the availability of initial spike sorting results for the given recording, containing manually verified single-unit clusters. In this way ground-truth data, which has both realistic spike and noise related statistics, can be made without modelling effort.

The input *high-pass filtered* extracellular recording that will serve as a basis for the hybrid ground-truth data is denoted by the matrix **X** ∈ ℝ^*T×N*^, with *T* the duration in samples of the recording and *N* the number of recording channels. The vector *x_c_* ∈ ℝ^*T*^ contains the elements from the *c*^th^ column of **X**, i.e., it contains the data recorded on channel *c*. A single sample from channel c at discrete time *k* is represented by x_*c*_ [*k*]. Let’s consider a single channel signal slice 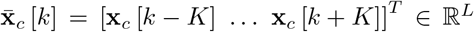 centered around *k*, where *L* = 2*K* +1 denotes the slice length, i.e., the so-called temporal window size in samples.

Initial spike sorting results are required for the estimation of the spike templates. The initial spike sorting results consist of a set of spike times 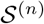 for every known single-unit cluster *n*. Consider the spike snippets matrix 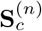 which contains in its columns all spike snippets 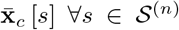. The spike template 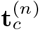 of a single-unit cluster n on channel c can then be estimated as follows:

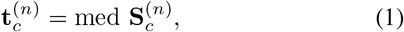

where the median operator acts on each row of the spike snippets matrix, such that 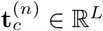.

Next, for every channel *c*, the template energy 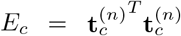 is calculated and the maximum energy *E*_max_ = max_*c*_ *E_c_* over all channels is determined. The channels in the template that contain relatively little energy, are considered to be noise channels and are zero-forced based on the following maximum energy criterion:

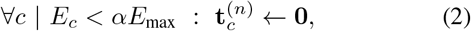

where *α* is the so-called zero force fraction (e.g., *α* = 0.03). This zero-forcing results in a more localized spike template.

Next, the optimal template scaling for every spike snippet used during the template estimation is calculated. This template scaling will be used both when subtracting and inserting units during the hybridization process. The optimal scaling is given by the least squares fitting factor 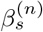 and minimizes the following least squares optimization problem:

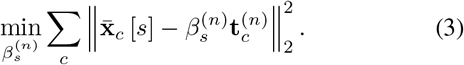

The solution to (3) is given by the following expression:

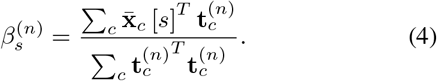

Before inserting the hybrid spikes into the recording, for every spike time used during the template estimation, the fitted template is subtracted from the data. Although this step is not strictly necessary, this subtraction will prevent the total energy in the final hybrid recording from increasing. The subtracted signal s_*c*_ [*k*] is given by:

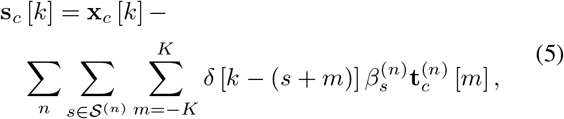

where *δ* [*k*] is the Kronecker delta, i.e., *δ* [*k*] = 1 if *k* = 0 and *δ* [*k*] = 0 if *k* ≠ 0.

The next step in the process of generating the hybrid ground truth is to change the spatial location of the spike template. In order to facilitate this spatial migration, some assumptions about the probe geometry have to be made. For every probe a rectangular grid model is assumed (non-rectangular probes are supported, but will be internally treated as rectangular grids, see below). The grid model requires that all recording electrodes for a certain probe lie on the intersection of an imaginary rectangular grid, as depicted in Fig. 1**A**. However, not every intersection point has to be occupied by an electrode, thereby allowing to model probes with broken or missing electrodes, or probes with a non-rectangular electrode grid. Furthermore, the model assumes that the horizontal distance between the intersection points is constant. Also the vertical distance between intersection points has to be constant, but this vertical distance can differ from the horizontal constant distance.

**Fig. 1.**
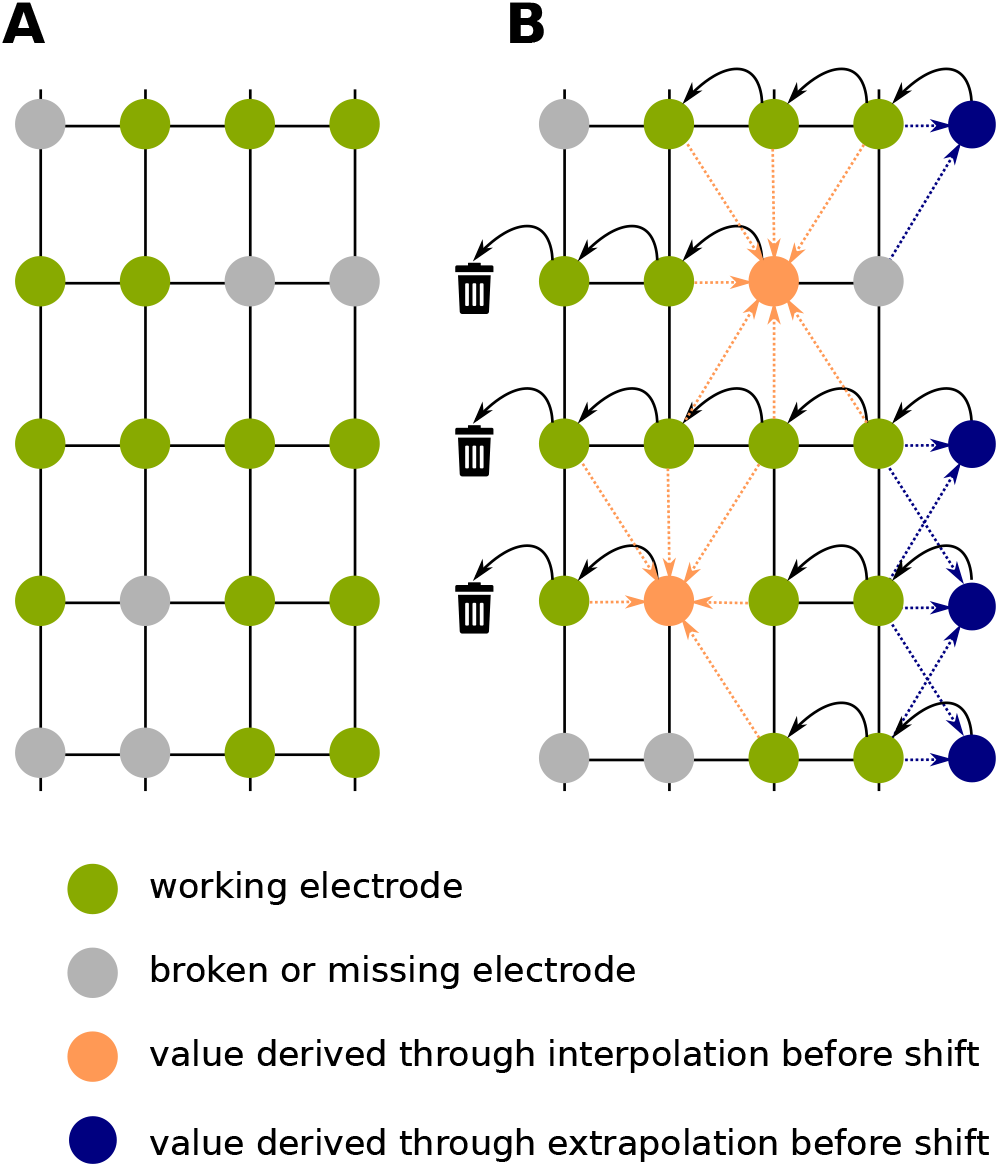
**A**: The probe grid model assumes that all electrodes (green dots) are located on intersections of a grid. The distance between grid lines along a certain axis is assumed to be constant. Not every intersection has to be populated by an electrode (grey dots). **B**: Moving the template over the probe consist of shifting the template over the grid. The black arrows represent shifting a template one position to the left. At the right side of the probe extrapolated waveforms (blue dots and arrows) are introduced into the template, while on the left side, a part of the template is lost. Missing channels that are shifted onto working electrodes contain a value obtained through interpolation (orange dots and arrows).

From the above assumptions a probe graph model can be built. For every empty location on the grid a missing electrode is added to the graph model. In such a graph model every electrode is aware of its neighbouring (missing) electrodes, such that the template can be easily shifted in every direction on the probe.

A template shift is characterized by a number of moves along the horizontal axis in a certain direction and a number of moves along the vertical axis in a certain direction. For the horizontal axis moving the template to the right is represented by a positive number and moving to the left by a negative number. Moving up is represented by a positive number of moves, while moving down has a negative sign. The shifted template 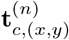 is obtained from shifting horizontally by x and shifting vertically by y. Fig. 1**B** gives a schematical representation of 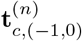 The black arrows in the figure indicate the spatial pseudo-permutation, note how extrapolated waveforms enter the shifted template (here by template we mean the set 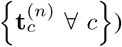 on the right side of the probe and how template information is dropped on the left side of the probe. Template extrapolation is performed to gracefully handle spikes at the edge of the probe, that might otherwise cause edge effect artefacts in case zero-padding is used rather than extrapolation. The extrapolated waveforms result from averaging over the neighbouring working electrodes (see blue discs in Fig. 1**B**) and an equal amount of fictitious electrodes containing all zeros to guarantee a major attenuation of the extrapolated waveforms when compared to their working neighbours. Missing channels that are shifted onto working electrodes contain a value obtained through interpolation (see orange discs in Fig. 1**B**). The interpolated value equals the average over all working neighbouring channels, as is schematically represented in Fig. 1**B** by the orange arrows.

Typically, the set of scaling parameters 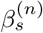 contains various outliers, e.g., caused by fitting the template on a segment that is deformed by the presence of overlapping spiking activity from other neurons. To prevent outliers in 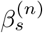 from propagating into the generated ground-truth data, only a subset of 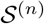 should be considered for reinsertion. This subset is derived by choosing a lower bound 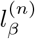 and upper bound 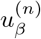 for the template scaling factor 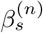 and setting 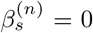 for all the spike times 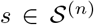 for which 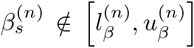, thereby effectively discarding those spikes from 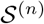.

Prior to the insertion of the ground truth spikes in the recording, a temporal within-sample-time jittering is applied to the template for every 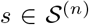. This jittering is obtained by first upsampling the template by a factor of 10, then applying a random within-sample-time shift sampled from a uniform distribution to the upsampled template, and finally, downsampling the template to the original sampling frequency. This temporal jittering models the fact that the occurrence of spikes are not phase locked with the sample clock of the analog-to-digital converter. The jitter operator is represented by the non-linear function *j_s_*: ℝ^L^ ↦ ℝ^*L*^, where the dependency on s is required to guarantee that for a certain spike time s all samples of the template are jittered by the same amount and because the random time shift is different for each 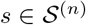. Finally, the hybrid data can be mathematically expressed as follows:g

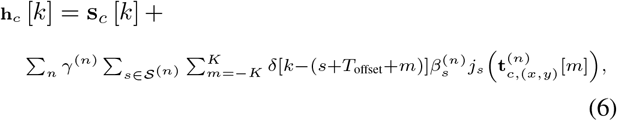

where we introduced an additional time offset *T*_offset_ to all remaining spike times before re-inserting them at the new location in order to decorrelate them with possible spike residues at the original location. *γ*^(*n*)^ is an optional user-defined scaling parameter to control the signal-to-noise-ratio (SNR) of the resulting hybrid unit associated with the neuron *n*. The default value is set to *γ*^(*n*)^ = 1.

## III. Gui for hybrid data generation

The generation of ground-truth data is a delicate process, which at least requires visual feedback on the generated ground truth and ideally some user interaction to make the key decisions during the generation process. For this reason we developed a Python-based GUI called SHYBRID (spike hybridizer) to support spike sorting users and developers with the generation of ground-truth data, so that they gain more insight into how spike sorting algorithms behave on certain data. This section elaborates on the typical usage flow of the tool. The following topics will be discussed in the different subsections: which input data to provide, how to visualize and adapt the template, why and how to choose bounds on template scaling, and what to consider when moving a unit across a probe. We will also touch on functionality to import and export templates for additional flexibility. Finally, we will discuss the tool’s functionality for the automatic assessment of spike sorting algorithms. This section only gives an overview of the above concepts, more detailed usage-related information can be found in the user manual that is provided with the tool.

### A. Input data

The user interface depends on four different inputs:

1. an extracellular multi-channel recording in binary format
2. the recording’s probe file
3. initial spike sorting results
4. a parameter file

The provided *binary recording* is assumed to be high-pass filtered. This high-pass filtering is standard practice in spike detection and spike sorting pipelines, as it is needed to attenuate the power dominant low frequency oscillations present in raw neural recordings. Without proper filtering, it would be difficult to reliably extract spike templates from the recording, which are the key building blocks for hybrid data generation. Only binary recordings that encode the signal samples using a signed data type are supported.

The application’s centerpiece is an intuitive data visualization widget. Signal snippets are plotted as if they are drawn on the probe (see Fig. 2). This visualization requires information about the geometry of the probe as described in the *probe file* [41] in a structured text format. This file format is used by some spike sorting algorithms already, so there are many probe files available for various neural recording devices. The probe file can also be used to inform the tool about broken/bad channels that are present in the recording, and are to be ignored during the hybridization process. Note that the tool is only compatible with single shank probe files. Some probe files also provide a probe graph. The SHYBRID will ignore any graph present in the probe file and will build its own. This graph is built from the geometry related grid assumptions as introduced in Section II.

**Fig. 2.**
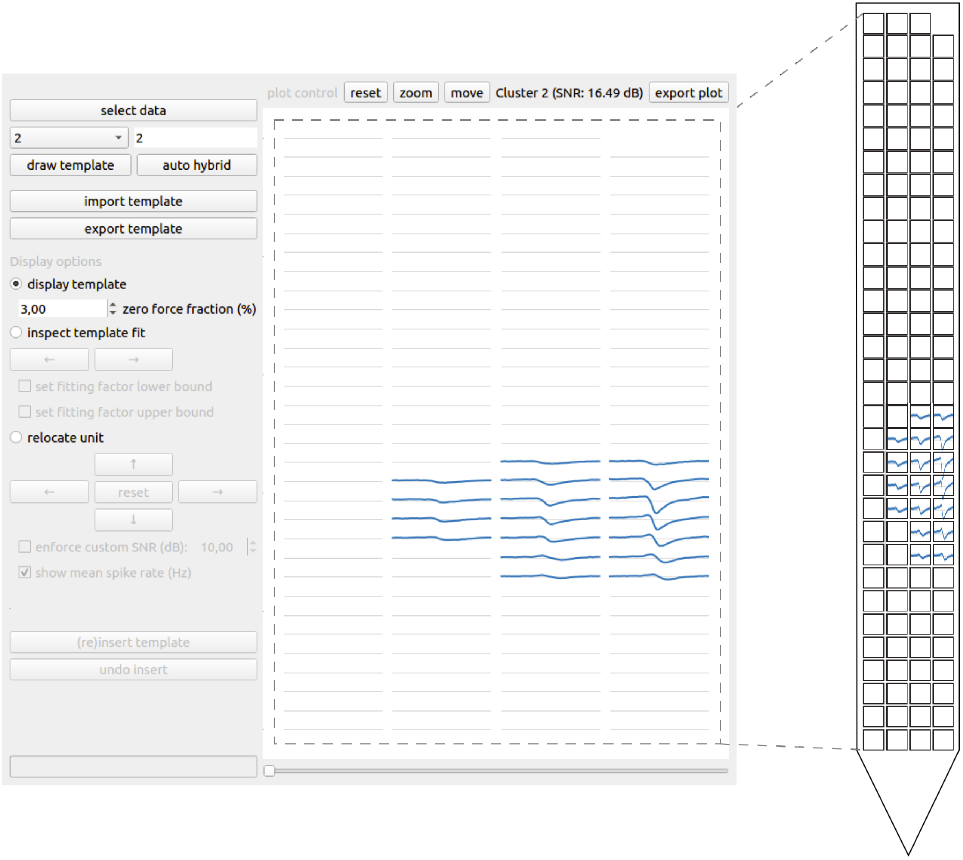
Spike templates (and recording snippets) are visually organized according to the channel geometry of the recording device. The different rows and columns in the visualization correspond to the rows and columns of the recording device. The waveform duration (horizontal axes) is controlled by the user. The waveform amplitude (vertical axes) is automatically scaled to provide a clean visualization. The relative waveform amplitude with respect to the recording noise is available from the SNR that is calculated and shown for every spike template.

The initial *spike sorting results* are a collection of manually curated single-unit clusters. A cluster consists of a unique integer identifier and the cluster’s corresponding spike times (given in samples). The initial spike sorting results can be loaded using either a custom *CSV*-file or by supplying a path to the folder containing the *phy* compatible data [41] in the *template-gui* format^2^ (i.e., a de facto standard for spike sorting results). The CSV-file has to consist of two columns, where the first column contains the cluster identifier and the second column contains a corresponding discrete spike time. From a CSV-file all clusters are loaded into the application. When using the phy compatible initial spike sorting results, only clusters that are labeled as “good” will be loaded.

Finally, the *parameter file* links all of the above files together as illustrated in Listing 1. On top of that, it contains additional recording-related parameters that are needed by the application, such as the recording’s sampling frequency and data type. The parameter file has to have the same file name as the binary recording file, and requires a *.yml* extension, indicating that its content is structured using the *YAML* format.

**Listing 1.**
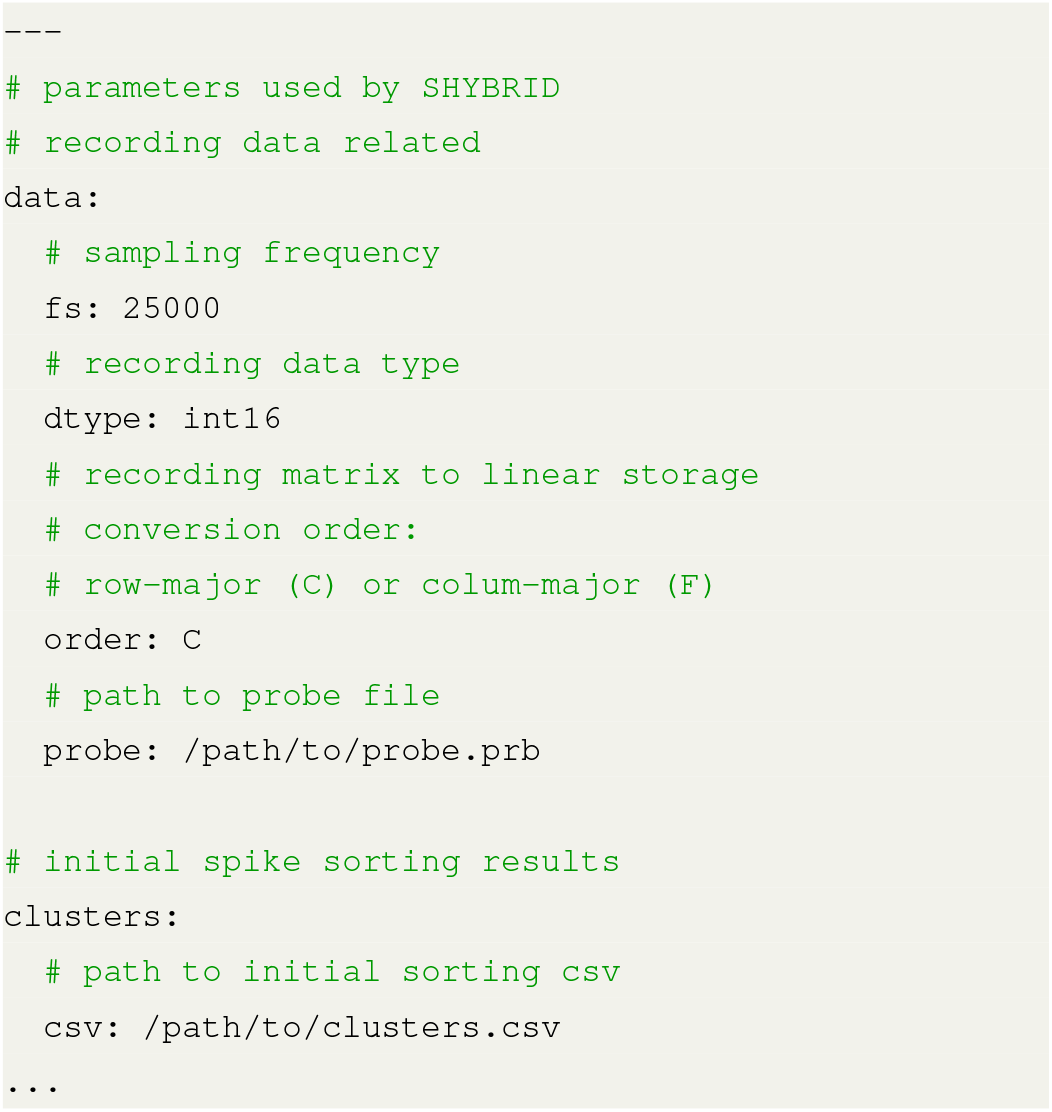
An example parameter file in the YAML format.

### B. Visualizing the template

After loading the input data, a spike template can be drawn for a single-unit spike cluster of choice. Details on how the template is estimated can be found in Section II. Two important parameters that affect the template estimate can be altered in the tool as shown in Fig. 3.

**Fig. 3.**
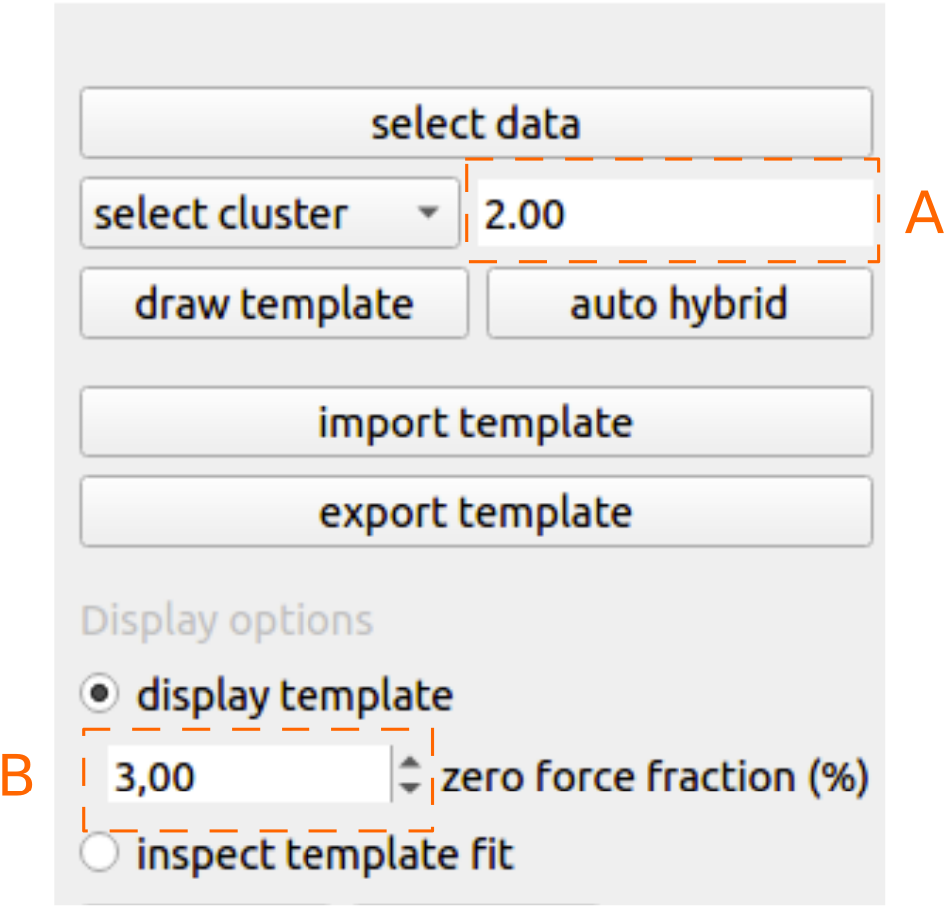
**A**: Template temporal window length given in milliseconds. **B**: Altering the zero force fraction changes the spatial extent of the template.

The first parameter that has to be explicitly chosen is the temporal window length. Changing this window length will change the number of consecutive samples on every channel around a single spike occurrence as used for template estimation and visualization. For generating hybrid data, this window length should be sufficiently large. Within the temporal window, the signal on every channel of the template should return to zero on both sides, this is to ensure that the complete spike template is caught in the given window. Choosing an appropriate window size will limit residual artefacts that occur during the process of hybrid data generation.

The second template related parameter that can be altered is the zero force fraction. Usually a spike template as seen on a probe is spatially sparse, i.e., some channels of the template are dominated by spiking activity, whereas others are dominated by noise. To eliminate any effect of these noise channels further down the pipeline, the tool tries to zero-force the noise channels. Typically these noisy channels are considerably lower in terms of signal energy. The software determines the signal energy for every channel of the template separately. From these energies the maximum energy is then determined. All channels that contain less energy than the given zero force fraction of the maximum template channel energy, are explicitly set to zero. As such the spatial extent of the template is determined from the data, instead of relying on an a-priori chosen spatial extent. The zero force fraction is set to 3% by default. Depending on the SNR of the recording, this fraction might have to be adjusted to reflect the true spatial footprint of the spike template. Increasing the zero force fraction (e.g., in case of a recording with low SNR) will eventually lead to more channels being set to zero.

After the template estimation process, the template is visualized as shown in Fig. 2. Only non-zero forced channels are emphasized (in blue) in the visualization.

### C. Inspecting template fit

A second step in the generation of hybrid data is to inspect how well a template fits the underlying data from which it was estimated. This is important to verify whether the scaled template model that is used for the generation of hybrid data is valid for the given cluster. For every data chunk that was used for the estimation of the template, an optimal template scaling factor is determined that minimizes the squared error between the data chunk and the scaled template (see Section II).

In the *inspect template fit* display mode, the scaled template (blue) is plotted on top of a signal chunk (red) containing a spike that was used during the template estimation process (see Fig. 4). By using the left and right arrow buttons (or the bottom slider) the user can browse through all the data chunks that were used for the estimation of the template. In this way the user can assess whether the scaled template sufficiently models the underlying data. To avoid that the user has to go exhaustively through all the chunks, the data chunk browsing order is determined by the fitting factor 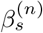, where moving the slider to the right corresponds to an increasing fitting factor.

The reason that we opted for this ordering is that often, both very low and high fitting factors are likely to be the reflection of a spurious template fit. Such a spurious template fit is typically caused by either false positives in the provided spike sorting results or are due to spatio-temporally overlapping spikes. Because of the ordering that is in use here, bounds can be easily identified and set on the fitting factors. The red profile at the bottom of the plotting area (see Fig. 4) is a visual representation of the ordered fitting factors for all data chunks. A user is adviced to start from the lowest scaling factor and move upwards until a nicely fitting signal chunk is identified. This chunk can then be set as the lower bound. After choosing a lower bound, the user can start from the maximum fitting factor and repeat the previous procedure in a downwards fashion. Only the spikes within the optional bounds will be considered during the hybridization process. This feature allows the user to have precise control over the scaling factor range, preventing unrealistic scaling factors from entering the hybrid data.

Because overlapping spikes will often cause unrealistic fitting factors, one might think that applying bounds on the fitting factors might exclude those interesting overlapping spike times from the hybridization process. However, since the spike template is spatially moved before the hybrid spikes are reinserted, it is very likely that new spatio-temporal overlap arises in the region to which the template is shifted.

### D. Relocating the unit

Once the validity of the scaled template model for the active cluster has been assessed, the unit has to be relocated on the probe. This action is fundamental for the generation of hybrid data, as mentioned before. Moving the unit along the probe also allows for the reintroduction of overlapping spikes, i.e., by moving the fictitious neuron to a location on the probe where there is a high spiking activity. To aid the user in choosing where to move the unit to, the average spike rate on every electrode is calculated and represented visually in the *relocate unit* view. By moving the unit to a *busy* region of the probe, it is more likely that spike overlap occurs for the hybrid neuron. If a unit is moved to a *silent* region, the hybrid spikes are more likely to be easily separable. This allows a user to control the difficulty level for the spike sorting algorithm under test. In our illustrative example, the unit that is shown in Fig. 4 is moved to a region with lower average spiking activity as shown in Fig. 5. The user can also control the difficulty level by enforcing a custom peak-SNR for the resulting hybrid unit. The peak-SNR is defined in Appendix C. By default, the peak-SNR is displayed corresponding to the default value *γ*^(*n*)^ = 1 in (6), which means the original amplitude of the spike template is kept.

**Fig. 4.**
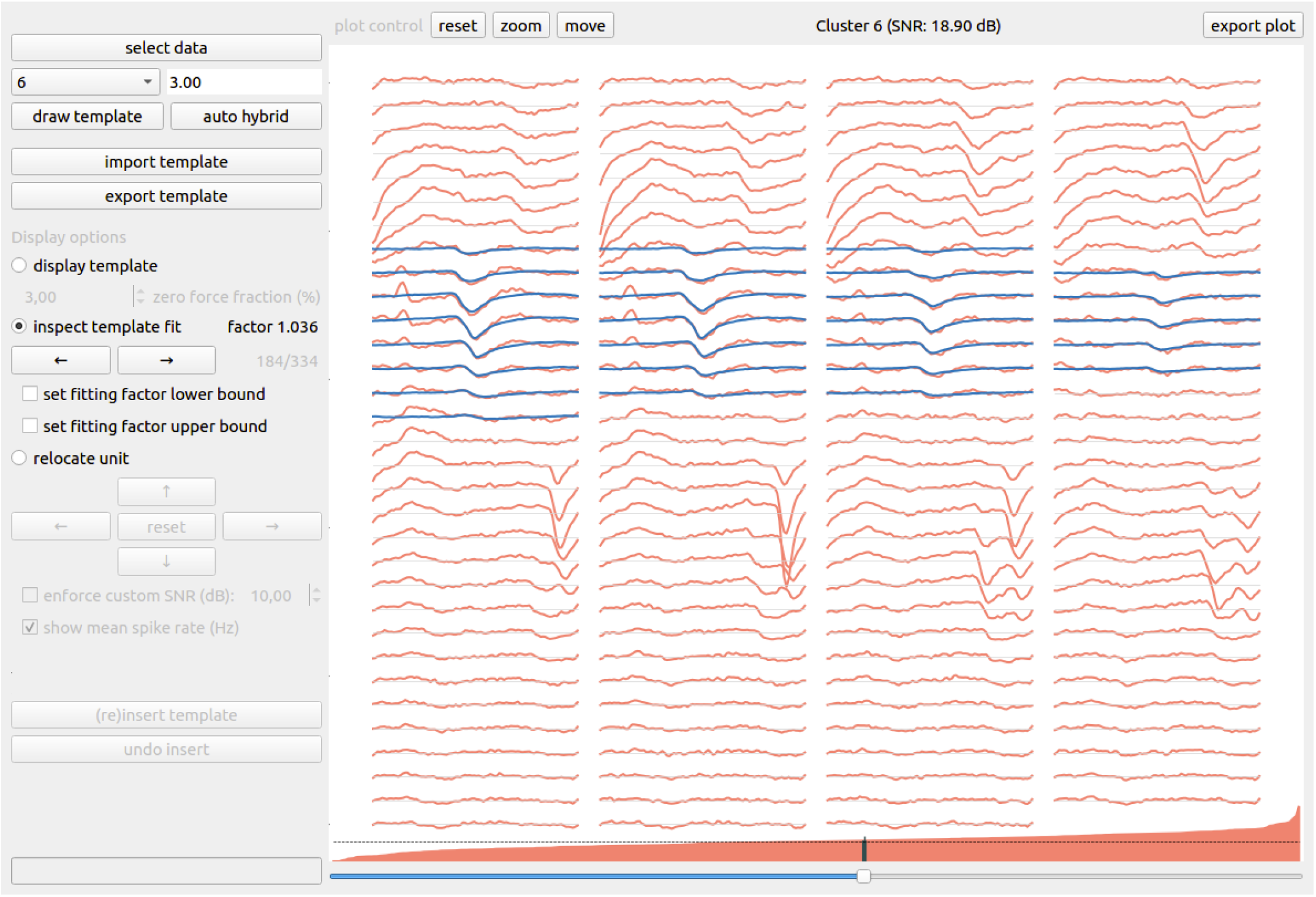
For every recording chunk (shown in red) that is used for the template estimation, an optimal template scaling factor is determined. The *inspect template fit* view allows the user to assess how good of a model the scaled template (shown in blue) is for the current cluster. The organization of the visualization is identical to Fig. 2. The red profile at the bottom of the plotting area is a visual representation of the ordered scaling factors for all recording chunks. The fine dashed line (black) indicates the unit fitting factor, i.e., 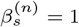.

**Fig. 5.**
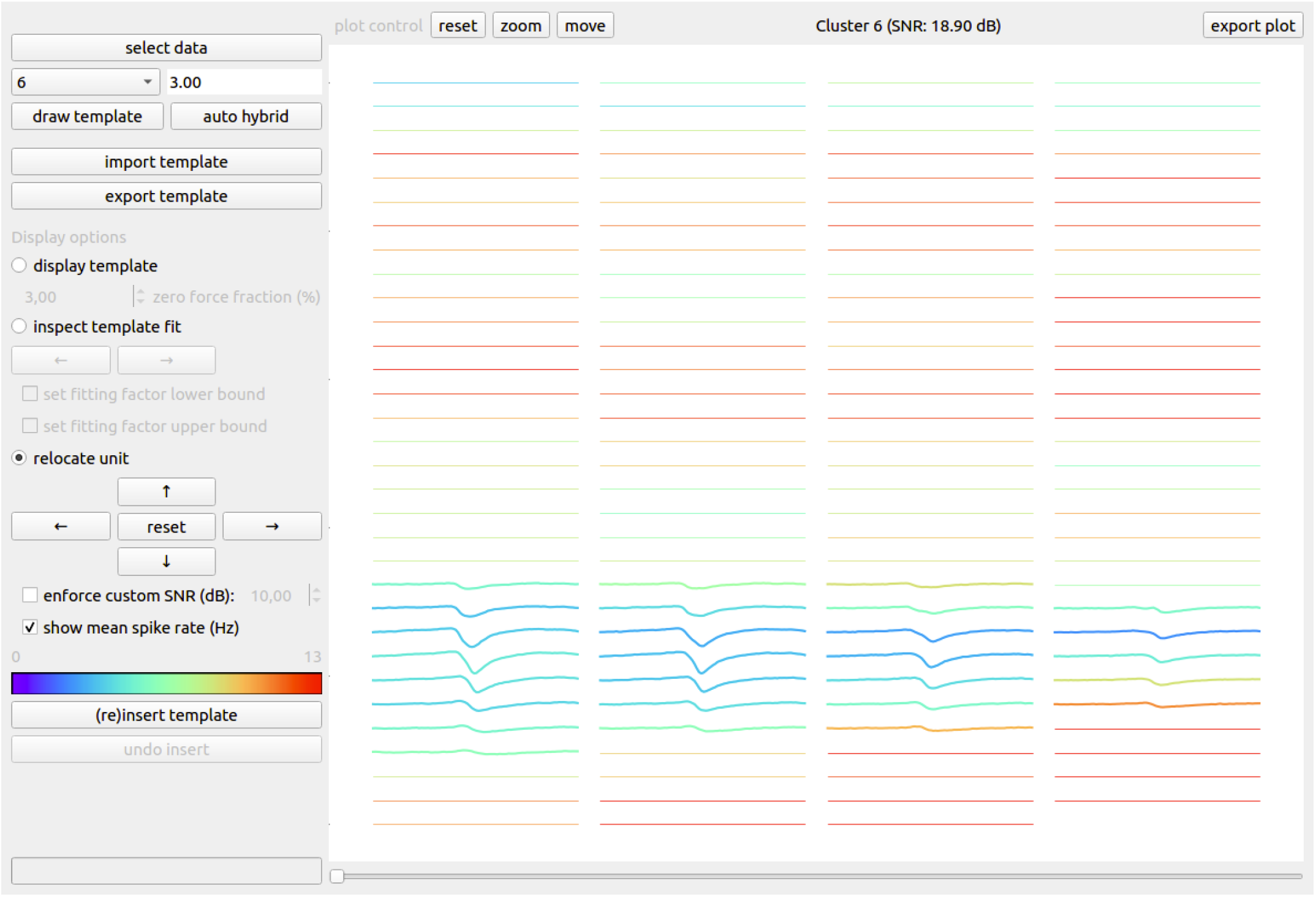
A unit’s spike template can be moved over the probe by using the arrow controls on the left panel. The average spike rate on every channel is indicated by the color in which the channel is plotted. Moving a template into a silent region will likely result in a hybrid recording that is easier to sort. The organization of the visualization is identical to Fig. 2.

After a unit is moved from its original spatial location, it has to be reinserted into its new location. A unit reinsertion first subtracts the original fitted templates from the recording after which the spikes within the template scaling factor bounds are reinserted at the new location. The spike times at which the scaled shifted templates are inserted are equal to the original spike times offset by an arbitrary constant that equals two times the temporal window length. As discussed in Section II, this offset is added to prevent the residual artefact after subtraction from interfering with the hybrid cluster template. This reinsertion will also create a CSV-file that keeps track of the hybrid ground-truth spike times. Note that the reinsertion is immediately applied to the provided recording and overwrites the binary data file. A reinsertion can always be undone if the resulting hybrid unit is not satisfactory, e.g., when the provided custom SNR is deemed unrealistic after inspecting the hybrid spike chunks.

### E. Auto hybridization

For long recordings containing dozens of manually curated single-unit clusters, the full user guided hybridization process is a lengthy procedure. Although we believe it is advisable to have a user making the key decisions, we also provide an auto hybridization function which loops over all provided single-unit clusters. For every cluster, conservative bounds on the template fitting factor are chosen and the corresponding unit is moved to another spatial location on the probe that is randomly chosen. All of this happens automatically, at the cost of reduced control on the resulting hybrid data. The details on the automatic bounds selection and template movement can be found in Appendix A and B, respectively.

### F. Importing / exporting templates

To provide increased flexibility it is also possible to import external spike templates into a recording. This feature allows for the creation of ground-truth data without the need for a prior spike sorting. The external templates can be either hand crafted or exported from other recordings that do have some prior spike sorting information that allows for the estimation of spike templates. Details about the importing procedure can be found in Appendix C.

### G. Automatic tools for spike sorting algorithm assessment

Since the main objective of this work is to get a better insight into spike sorting performance, the tool comes with methods for comparing the hybrid data spike sorting results to the hybrid ground-truth spike times. This functionality consists of two important methods, that will be discussed in more detail starting from the next paragraph. First, the functionality to perform automatic cluster merging on the spike sorting results is discussed. Second, the spike sorting performance metrics routine is discussed.

#### 1) Automatic cluster merging

Ideally, a spike sorting algorithm outputs a single cluster for each single-unit spike train present in a recording. In practice however, spikes from the same neuron are often split over multiple clusters (referred to as overclustering) or a single cluster contains spikes from multiple neurons. Experimentalists usually prefer overclustering, because the manual curation which then consist of cluster merging is more straightforward than the process of splitting clusters into several single-unit clusters. Therefore, modern spike sorting algorithms are typically tuned towards overclustering.

If an algorithm under investigation is known to have a preference for overclustering, it is necessary to perform a merging step prior to calculating the spike sorting performance metrics. For this purpose an automatic ground truth assisted merging framework is implemented to bypass this manual merging step in the assessment of a spike sorting algorithm. Since we do not provide an automatic splitting framework, algorithms that are tuned towards overclustering are favoured in terms of final spike sorting accuracy. We believe this limitation is acceptable, as it reflects the user preference towards overclustering. The algorithmic details of the merging framework are given in Appendix D.

#### 2) Performance metrics calculation

A fast implementation is provided for the calculation of spike sorting related performance metrics given the hybrid ground-truth data. The provided method is capable of generating tables such as Table I and Table II. Prior to the calculation of the metrics, for every hybrid spike train an automatic cluster merging step is performed as explained above. After this merging step, the following metrics are calculated for every ground-truth spike train:

- The *recall* or sensitivity, which is defined as 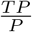, with TP the number of true positives, i.e., the number of spikes that are correctly associated with the ground-truth spike train and P the number of positives, i.e., the total number of spikes in the ground-truth spike train.
- The *precision,* which is defined as 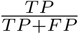, with FP the false positives, i.e, spikes that are wrongly associated with the ground-truth spike train.
- The F_1_-score is also calculated, which is the harmonic average of the recall and precision: 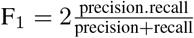.
- The number of clusters that were automatically merged, prior to calculating the above performance metrics.

**Table I.**
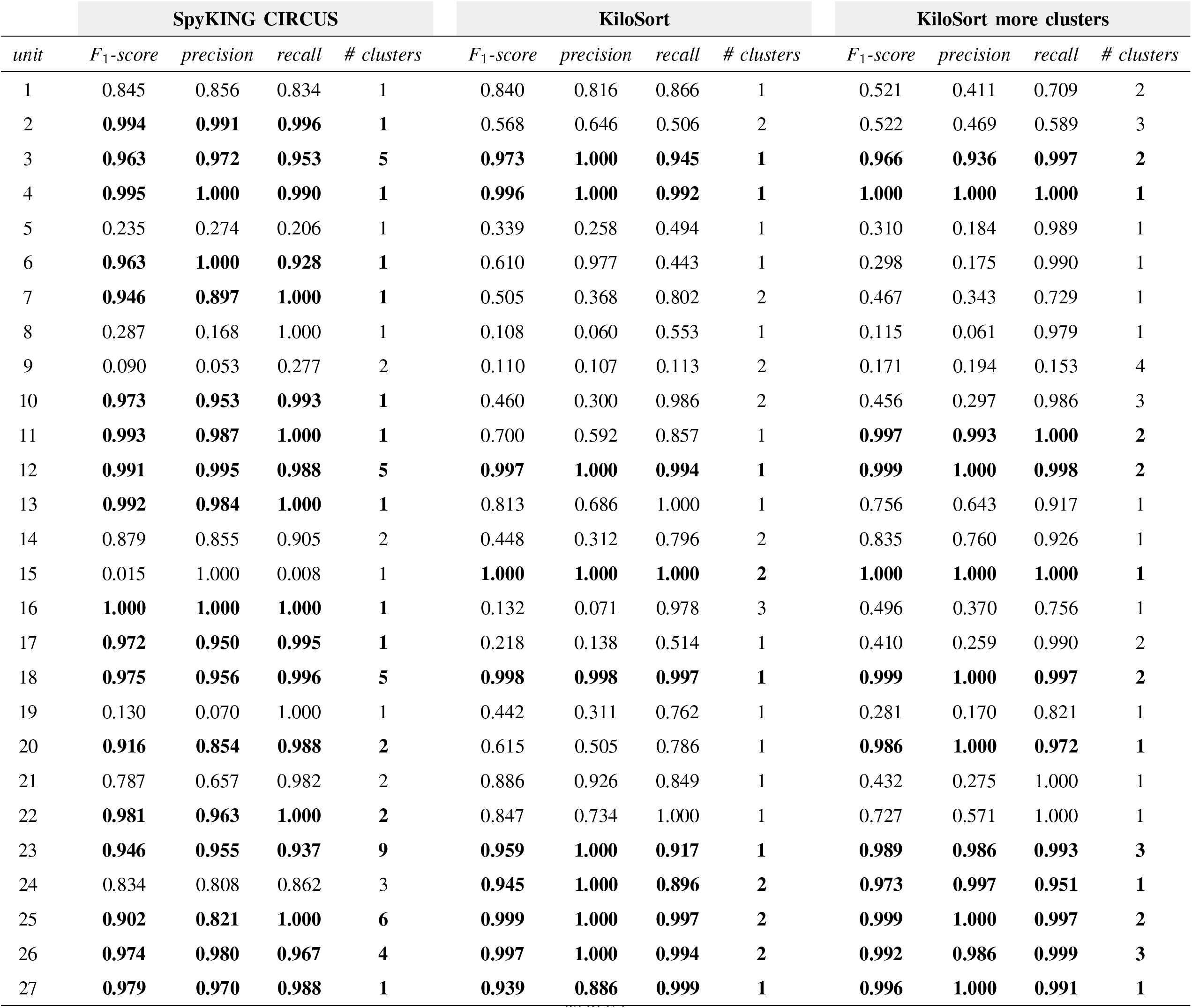
Spike sorting performance metrics for the three separate spike sorting runs (SC, KS, KS more clusters) applied on the SC-informed hybrid ground-truth data. For every ground-truth unit in every spike sorting run, the F_1_-score, precision, recall and number of cluster merges is shown. Rows with bold typesetting indicate the ground-truth units that were recovered as single-unit spike trains. This table is generated by shybrid.

| SpyKING CIRCUS |  |  |  |  | KiloSort |  |  |  | KiloSort more clusters |  |  |  |
| --- | --- | --- | --- | --- | --- | --- | --- | --- | --- | --- | --- | --- |
| unit | $F_1$-score | precision | recall | # clusters | $F_1$-score | precision | recall | # clusters | $F_1$-score | precision | recall | # clusters |
| 1 | 0.845 | 0.856 | 0.834 | 1 | 0.840 | 0.816 | 0.866 | 1 | 0.521 | 0.411 | 0.709 | 2 |
| 2 | **0.994** | **0.991** | **0.996** | **1** | 0.568 | 0.646 | 0.506 | 2 | 0.522 | 0.469 | 0.589 | 3 |
| 3 | **0.963** | **0.972** | **0.953** | **5** | **0.973** | **1.000** | **0.945** | **1** | **0.966** | **0.936** | **0.997** | **2** |
| 4 | **0.995** | **1.000** | **0.990** | **1** | **0.996** | **1.000** | **0.992** | **1** | **1.000** | **1.000** | **1.000** | **1** |
| 5 | 0.235 | 0.274 | 0.206 | 1 | 0.339 | 0.258 | 0.494 | 1 | 0.310 | 0.184 | 0.989 | 1 |
| 6 | **0.963** | **1.000** | **0.928** | **1** | 0.610 | 0.977 | 0.443 | 1 | 0.298 | 0.175 | 0.990 | 1 |
| 7 | **0.946** | **0.897** | **1.000** | **1** | 0.505 | 0.368 | 0.802 | 2 | 0.467 | 0.343 | 0.729 | 1 |
| 8 | 0.287 | 0.168 | 1.000 | 1 | 0.108 | 0.060 | 0.553 | 1 | 0.115 | 0.061 | 0.979 | 1 |
| 9 | 0.090 | 0.053 | 0.277 | 2 | 0.110 | 0.107 | 0.113 | 2 | 0.171 | 0.194 | 0.153 | 4 |
| 10 | **0.973** | **0.953** | **0.993** | **1** | 0.460 | 0.300 | 0.986 | 2 | 0.456 | 0.297 | 0.986 | 3 |
| 11 | **0.993** | **0.987** | **1.000** | **1** | 0.700 | 0.592 | 0.857 | 1 | **0.997** | **0.993** | **1.000** | **2** |
| 12 | **0.991** | **0.995** | **0.988** | **5** | **0.997** | **1.000** | **0.994** | **1** | **0.999** | **1.000** | **0.998** | **2** |
| 13 | **0.992** | **0.984** | **1.000** | **1** | 0.813 | 0.686 | 1.000 | 1 | 0.756 | 0.643 | 0.917 | 1 |
| 14 | 0.879 | 0.855 | 0.905 | 2 | 0.448 | 0.312 | 0.796 | 2 | 0.835 | 0.760 | 0.926 | 1 |
| 15 | 0.015 | 1.000 | 0.008 | 1 | **1.000** | **1.000** | **1.000** | **2** | **1.000** | **1.000** | **1.000** | **1** |
| 16 | **1.000** | **1.000** | **1.000** | **1** | 0.132 | 0.071 | 0.978 | 3 | 0.496 | 0.370 | 0.756 | 1 |
| 17 | **0.972** | **0.950** | **0.995** | **1** | 0.218 | 0.138 | 0.514 | 1 | 0.410 | 0.259 | 0.990 | 2 |
| 18 | **0.975** | **0.956** | **0.996** | **5** | **0.998** | **0.998** | **0.997** | **1** | **0.999** | **1.000** | **0.997** | **2** |
| 19 | 0.130 | 0.070 | 1.000 | 1 | 0.442 | 0.311 | 0.762 | 1 | 0.281 | 0.170 | 0.821 | 1 |
| 20 | **0.916** | **0.854** | **0.988** | **2** | 0.615 | 0.505 | 0.786 | 1 | **0.986** | **1.000** | **0.972** | **1** |
| 21 | 0.787 | 0.657 | 0.982 | 2 | 0.886 | 0.926 | 0.849 | 1 | 0.432 | 0.275 | 1.000 | 1 |
| 22 | **0.981** | **0.963** | **1.000** | **2** | 0.847 | 0.734 | 1.000 | 1 | 0.727 | 0.571 | 1.000 | 1 |
| 23 | **0.946** | **0.955** | **0.937** | **9** | **0.959** | **1.000** | **0.917** | **1** | **0.989** | **0.986** | **0.993** | **3** |
| 24 | 0.834 | 0.808 | 0.862 | 3 | **0.945** | **1.000** | **0.896** | **2** | **0.973** | **0.997** | **0.951** | **1** |
| 25 | **0.902** | **0.821** | **1.000** | **6** | **0.999** | **1.000** | **0.997** | **2** | **0.999** | **1.000** | **0.997** | **2** |
| 26 | **0.974** | **0.980** | **0.967** | **4** | **0.997** | **1.000** | **0.994** | **2** | **0.992** | **0.986** | **0.999** | **3** |
| 27 | **0.979** | **0.970** | **0.988** | **1** | **0.939** | **0.886** | **0.999** | **1** | **0.996** | **1.000** | **0.991** | **1** |

**Table II.** Spike sorting performance metrics for the three separate spike sorting runs (SC, KS, KS more clusters) applied on the KS-informed hybrid ground-truth data. For every ground-truth unit in every spike sorting run, the F_1_-score, precision, recall and number of cluster merges is shown. Rows with bold typesetting indicate the ground-truth units that were recovered as single-unit spike trains. This table is generated by shybrid.

| SpyKING CIRCUS |  |  |  |  | KiloSort |  |  |  | KiloSort more clusters |  |  |  |
| --- | --- | --- | --- | --- | --- | --- | --- | --- | --- | --- | --- | --- |
| unit | F<sub>1</sub>-score | precision | recall | # clusters | F<sub>1</sub>-score | precision | recall | # clusters | F<sub>1</sub>-score | precision | recall | # clusters |
| 1 | 0.298 | 0.192 | 0.660 | 1 | 0.106 | 0.056 | 0.992 | 3 | 0.139 | 0.139 | 0.140 | 3 |
| 2 | 0.678 | 0.516 | 0.988 | 1 | 0.643 | 0.474 | 1.000 | 2 | 0.601 | 0.430 | 1.000 | 3 |
| 3 | 0.283 | 0.165 | 0.998 | 1 | 0.702 | 0.546 | 0.984 | 1 | 0.440 | 0.415 | 0.469 | 2 |
| 4 | 0.084 | 0.070 | 0.105 | 8 | 0.047 | 0.043 | 0.053 | 13 | 0.090 | 0.065 | 0.143 | 29 |
| 5 | **0.928** | **0.915** | **0.941** | **4** | 0.893 | 0.991 | 0.813 | 1 | 0.364 | 0.263 | 0.590 | 9 |
| 6 | **0.936** | **0.928** | **0.943** | **8** | **0.962** | **0.976** | **0.948** | **2** | **0.983** | **0.981** | **0.985** | **5** |
| 7 | **0.952** | **0.911** | **0.997** | **1** | 0.263 | 0.154 | 0.903 | 3 | 0.629 | 0.481 | 0.909 | 4 |
| 8 | **0.999** | **0.998** | **1.000** | **3** | **0.999** | **1.000** | **0.998** | **2** | **0.988** | **0.978** | **0.998** | **1** |
| 9 | 0.433 | 0.337 | 0.604 | 2 | 0.789 | 1.000 | 0.652 | 2 | 0.613 | 0.622 | 0.604 | 4 |
| 10 | **0.962** | **0.974** | **0.951** | **6** | **0.999** | **0.999** | **0.999** | **2** | **0.997** | **0.994** | **0.999** | **2** |
| 11 | 0.808 | 0.971 | 0.693 | 2 | 0.633 | 0.465 | 0.991 | 1 | 0.631 | 0.537 | 0.764 | 1 |
| 12 | 0.615 | 0.481 | 0.854 | 2 | 0.685 | 0.527 | 0.978 | 1 | 0.683 | 0.519 | 1.000 | 2 |
| 13 | **0.920** | **0.869** | **0.978** | **6** | **0.907** | **0.832** | **0.997** | **3** | 0.866 | 0.965 | 0.785 | 1 |
| 14 | 0.800 | 0.993 | 0.670 | 1 | **0.997** | **0.997** | **0.997** | **2** | **0.980** | **1.000** | **0.961** | **4** |
| 15 | **0.971** | **0.983** | **0.960** | **6** | 0.702 | 0.542 | 0.997 | 1 | **0.972** | **1.000** | **0.945** | **3** |

The implementation is based on Python set structures. A spike train is modelled as a set containing the spike times at which the underlying neuron is active. A Python set is essentially a hash table. A hash table allows to check whether an element is present in the structure in O(1), i.e., the time it takes is independent of the number of spike times that are present in the set. This particular implementation based on hash tables allows for a very fast calculation of the above metrics. To cope with spike time misalignment between the ground truth and the actual spike sorting results, every spike time in the set structure is extended by a user-defined window.

### H. SpikeInterface integration

SpikeInterface [42] is an open-source software stack, that was designed to promote the interoperability between different neural recording systems and spike sorting software. This goal is reached by implementing a unified access model for both neural recordings and spike sorting. SpikeInterface also provides functionality for ground truth validation and curation that can be applied to their unified data access model. In order to further improve the user experience of SHYBRID, a SHYBRID-SpikeInterface integration is provided. This integration has the following advantages:

- Extend the native file format compatibility of SHYBRID with the file formats that are supported in the SpikeInterface ecosystem. This is both applicable to the neural recordings, as well as the initial spike sorting format.
- SHYBRID hybrid data can be easily sorted using the various spike sorters that are supported in the SpikeInterface ecosystem.
- Besides the SHYBRID validation routine, one can also analyse the hybrid data spike sorting results and perform a ground truth validation using SpikeInterface.

## IV. Case study: comparing spike sorting algorithms and tuning parameters

In this section the performance of two spike sorting algorithms on a specific recording is analysed. The two spike sorting algorithms considered here are SpyKING CIRCUS (SC) [18] and KiloSort (KS) [15]. The donor recording used here is part of a paired recordings dataset (2015 0903 CelL9.0) [24], which is commonly used for validating spike sorting algorithms [17] [18] [19].

The donor recording is given to both spike sorting algorithms prior to the hybridization process. We generate two hybrid data sets from this donor recording, where one is informed by the SC spike sorting (with default parameter settings) results and the other by the KS spike sorting (with default parameter settings, where *number of clusters* = 256) results. This will allow us later on to identify a potential bias towards the algorithm from which the spike sorting results are used during the hybridization process. The spike sorting results from both algorithms are manually curated using the phy template GUI [41]. This curation process consists of a manual cluster merging and assessing whether or not the cluster consist of single-unit activity. Two hybrid data sets are then generated, following the steps described in Section III, from the single-unit spike sorting results obtained during the manual curation of both algorithms. The total number of injected hybrid ground-truth units is 27 for the SC-informed hybrid data and 15 for the KS-informed hybrid data. There are no overlapping spike trains between the SC-informed and KS-informed data. Fig. 6 shows the spike templates of four hybrid units. This figure contains templates from both SC-informed and KS-informed hybrid data. Visual comparison shows that units that are easily recovered (see next paragraph for the definition) during the final spike sorting, are likely to have a higher SNR compared to units that were not recovered during this spike sorting.

**Fig. 6.**
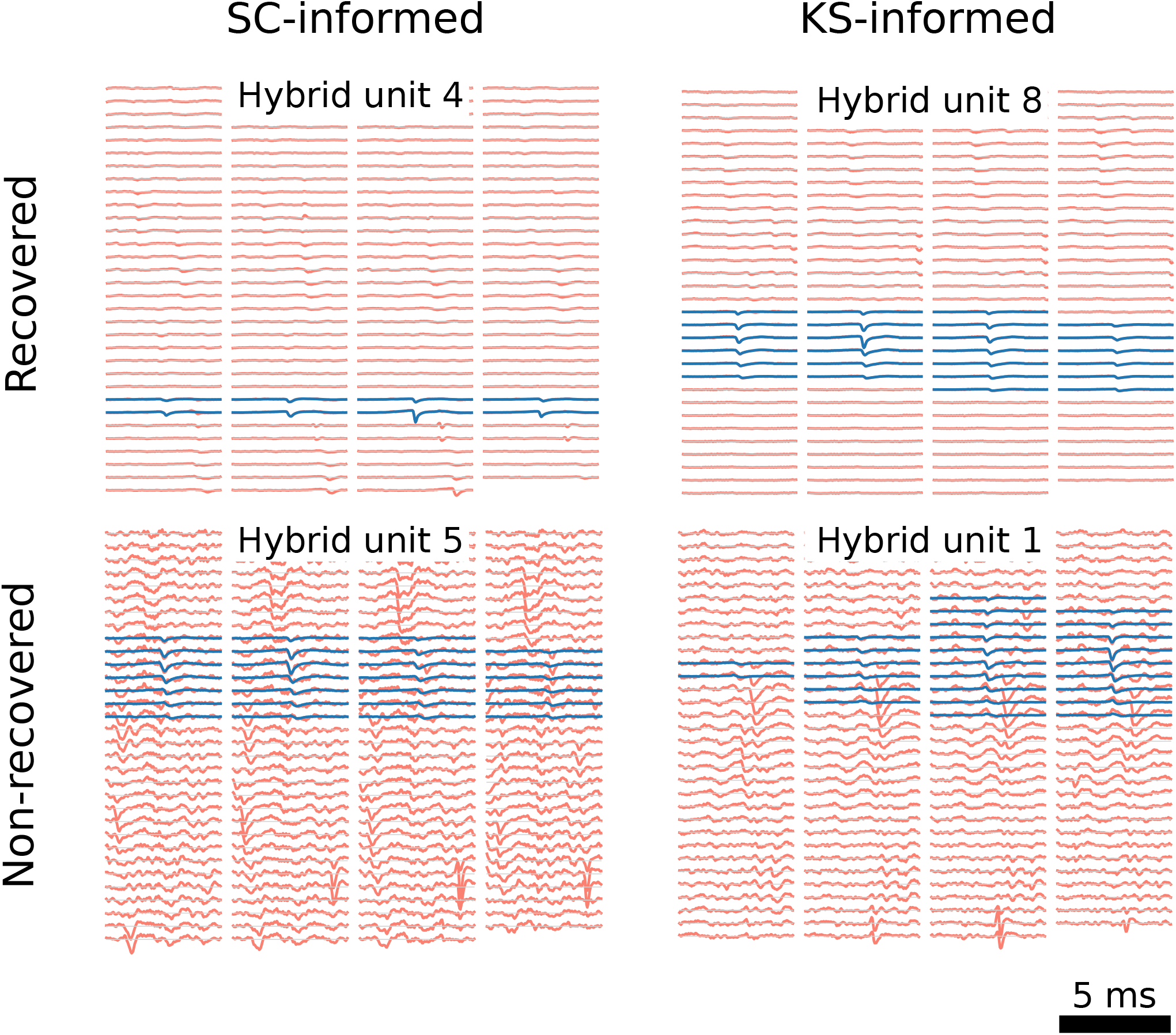
Example templates (blue) of hybrid units for both the SC-informed and KS-informed hybrid data are shown. The two templates in the top row originate from hybrid units that are recovered (i.e., F_1_-score > 0.9) by both spike sorting algorithms (see TABLE I and II for numerical details). The bottom row contains two templates from hybrid units that were only partially recovered by both algorithms. An example optimally scaled spike signal chunk (red) is shown behind each template to allow for a visual assessment of the relative noise levels with respect to the template power. The individual unit template plots are generated through the SHYBRID *export plot* functionality.

After hybridization, both hybrid data sets are again sorted using both algorithms (again with default parameter settings). Because this time the spike sorting is performed on groundtruth data, a ground truth assisted automatic merging can be performed after the spike sorting, as explained in Section III-G1. These automatically merged spike sorting results are then compared to the ground-truth labels. The automatic merging and ground truth comparison result in Table I and II. The rows in those tables that have a bold typesetting indicate ground-truth units that have been properly recovered, i.e., the F_1_-score for those units is greater than 0.9. Although this definition is somewhat arbitrary, it allows us to focus on the spike sorting performance metrics for units that have been successfully recovered.

The spike sorting performance metrics for the recovered units are also summarized in Fig. 7. Fig. 7**A** shows that for the SC-informed hybrid data SC recovers 18/27 units (dark blue bar) and KS recovers 10/27 units (red bar). One can also see that for the KS-informed hybrid data SC recovers 7/15 units and KS recovers 5/15 units. For both the SC and KS-informed hybrid data, the average number of automatic cluster merges is higher for SC than for KS (see Fig. 7**B**), and the average F_1_-score is higher for KS (red bar) than for SC (dark blue bar) as can be seen from Fig. 7**C** and 7**D**.

**Fig. 7.**
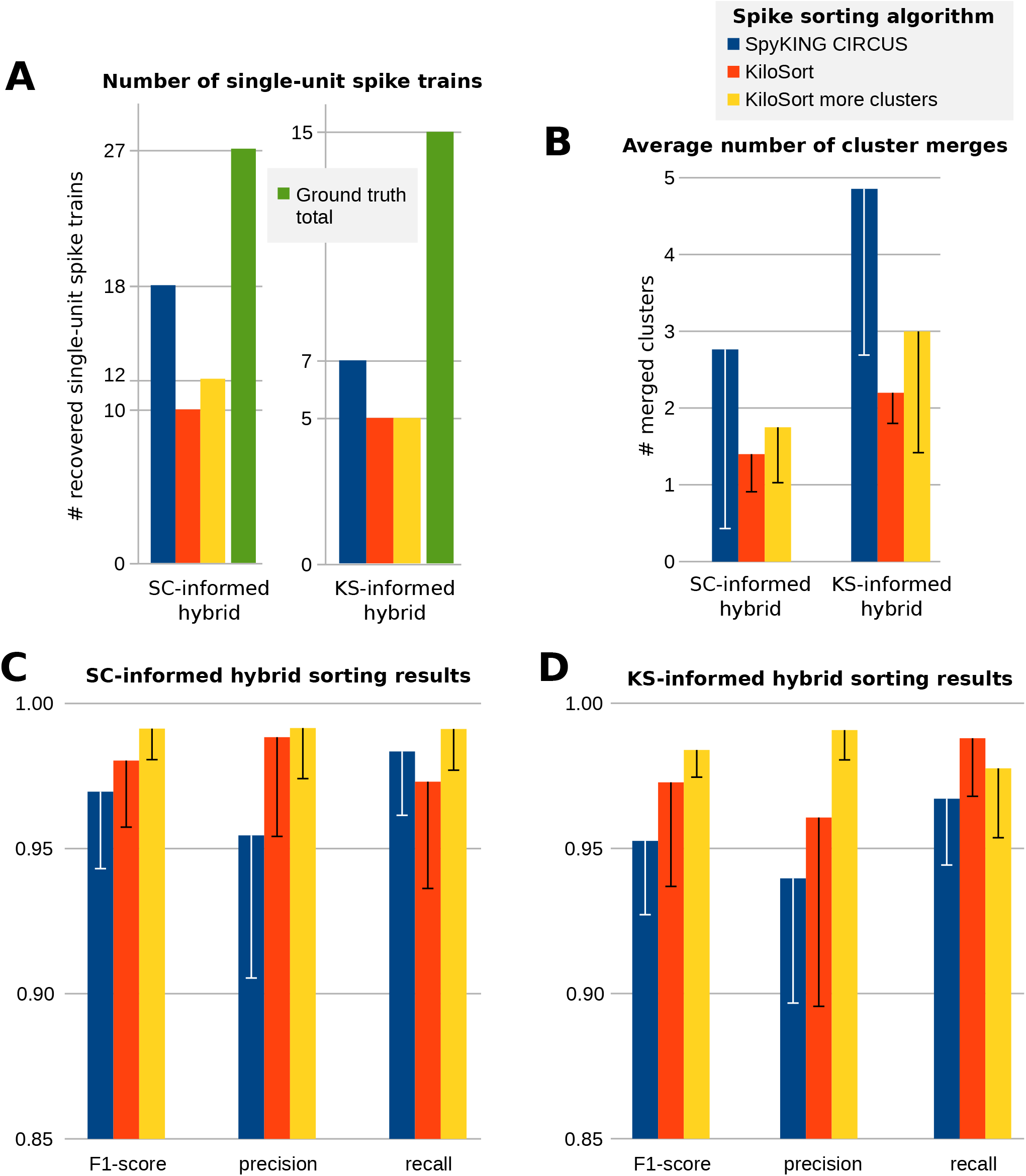
**A**: For both the SC- and KS-informed hybrid ground-truth data, the number of hybrid single-unit spike trains that are recovered by the different spike sorting algorithms is shown. The total number of hybrid ground-truth spike trains, i.e, the maximum number that could have been recovered, is shown in grey. **B**: For both the SC-and KS-informed hybrid ground-truth data, the average number of cluster merges over the single-unit spike trains is shown for the different spike sorting algorithms that are used. The error bars indicate the standard deviation. **C** and **D**: Spike sorting performance metrics for the SC- and KS-informed hybrid data, respectively. As performance metrics the F_1_-score, precision and recall are shown. The error bars indicate the standard deviation across the recovered single-unit spike trains.

SC recovers 66.7% of the hybrid units for the SC-informed data and 46.7% of hybrid units for the KS-informed data. The average F_1_-score for SC on the SC-informed data equals 97.0%, and 95.3% on the KS-informed data. This information might indicate the existence of a bias towards the initial spike sorter, when SC is used for that purpose. On the other hand, KS recovers 37.0% of the hybrid units for the SC-informed data and 33.3% for the KS-informed data. The average F_1_-score for KS on the SC-informed data equals 98.0%, and 97.2% on the KS-informed data. From this information we cannot conclude the existence of a bias towards the initial spike sorter, when KS is used for this purpose. Apart from the potential existence of a bias, the above results might also be partially explained by differences in spike sorting difficulty between the two different hybrid data sets.

From these first observations we can conclude that SC returns more single-unit clusters compared to KS for this specific data. However, the spike sorting accuracy is higher for the single-unit clusters retrieved by KS. The average KS unit also consists of fewer merged sub-clusters, indicating that less effort has to be spend on the manual curation. Based on these identified differences, we can differentiate between the two algorithms for this specific recording setting and choice of spike sorting parameters depending on the intended further use of the spike sorting results. In case these spike sorting results are further used for the estimation of neural population dynamics, we argue that having a higher number of single units at a slightly reduced spike sorting performance is favourable (motivated by [43]), as such favouring SC for this application. However, when these spike sorting results are used for investigating the exact role of a single neuron, e.g., investigating its spatial tuning [6], the quality of the sorting is more important than the quantity of retrieved single units. As such, favouring KS over SC in this context. More generally speaking, depending on which sorting characteristics are favourable for a specific application, an informed choice can be made between spike sorting algorithms by using hybrid data.

A key difference between both spike sorting algorithms compared here, is that KS requires the user to define the number of clusters prior to the actual spike sorting. The KS spike sorting results discussed above, used a value of 256 for the number of clusters. To try to increase the number of recovered single-unit spike trains for KS, the number of clusters is chosen as 512, while keeping the default setting for all other parameters. The results of this second KS run are represented in Fig. 7 by the yellow bars.

As can be seen from Fig. 7**A**, there is no direct evidence that KS with more clusters recovers more single-unit spike trains. only two additional single-unit spike trains are recovered under the new number of clusters setting for the SC-informed data. However, the F_1_-score for the recovered single-unit spike trains does increase even further, as compared to the previous spike sorting runs, as can be seen from Fig. 7**C** and 7**D**. As can be seen from this example, it is non-trivial to predict the effect of changing spike sorting related parameters on the spike sorting performance.

## V. Discussion and Conclusion

In this work we first formally introduced the hybrid groundtruth model. We presented a graphical tool to aid the creation of such hybrid ground-truth data. This graphical spike hybridizer for extracellular recordings or SHYBRID makes the creation of hybrid data very accessible to experimenters and tries to improve a user’s spike sorting experience. Because of the visual approach, a user can easily verify the key hybrid model assumptions and monitor the quality of the resulting hybrid recordings. Besides the data generation aspect, functionality is provided for automatic cluster merging and very time-efficient spike sorting performance metric calculation through hash functions.

A case study was conducted where the performance of two different spike sorting algorithms on a hybridized recording was investigated. From the spike sorting results obtained on the hybrid ground truth, clear differences in spike sorting characteristics between the two algorithms could be identified for this data. Such objectively identified differences can help a user to make an informed decision about which algorithm to use for a certain application, depending on the targeted algorithmic performance characteristics. During the case study the effect of changing spike sorting related parameters was investigated. It was demonstrated that the effect of adapting a parameter that controls the total number of spike clusters does not necessarily have the anticipated effect of recovering more single-unit spike trains. Hybrid ground-truth data allows for an objective quantification of the effect of specific parameter settings. By studying the effect of different parameter settings, a deeper understanding of spike sorting algorithms is promoted.

The use of hybrid ground-truth data should lead to a higher spike sorting performance and less time spent on manual curation when applied to similar future data. This is especially interesting for chronic recordings, where the measurement equipment is kept in the same brain region over multiple weeks to months. In such experiments a multitude of recordings is available from the same region. Especially for such chronic experiments, the initial hybridization effort might be worth the investment. It is needless to say that this tool is also valuable for the community of spike sorting developers. By sharing generated hybrid data within the community, a large body of extracellular ground-truth data can be obtained for future benchmarks.

Almost simultaneously with the release of SHYBRID, the MEArec [31] testbench simulator for ground-truth extracellular spiking data was released. The development of this tool is illustrative for the need of spiking ground-truth data for validating and better understanding spike sorting algorithms. MEArec generates its ground-truth data from computational simulations, using complex biophysically detailed models. Such a modelling approach enables full control over complex phenomena such as, e.g., bursting activity, drift or spatiotemporal synchrony. However, this high level of control implies that users have to acquire expert technical know-how in order to generate meaningful data, in particular when the aim is to generate ground-truth data that is representative for a specific experiment and/or recording set-up. Therefore, MEArec seems mostly suited for, e.g., spike sorting developers or modelling experts rather than spike sorting users.

SHYBRID does not require such a deep level of technical modelling expertise to generate meaningful ground-truth data. Furthermore, the ground-truth data is fully generated from an actual recording, which makes it intrinsically tailored to the recording setting of the user. The ease of use of SHYBRID comes at the expense of reduced control over certain recording-related characteristics, as compared to MEArec. Bursting cells can be included in the hybrid ground truth data, if they are present in the prior spike sorting results. Note that such SHYBRID bursting cells are only a simple amplitude modulation model of bursting, and do not account for template shape modulation that might occur for true bursting cells. Spatio-temporal overlap arises naturally when relocating units during the hybridization process, but synchrony can not be enforced. Drift simulation is not supported in SHYBRID for the time being, because it does not arise naturally from the presented hybrid model. A drift simulation framework could, technically speaking, be added to our software in a future update, but this will come at the cost of increased complexity for the user. SHYBRID and MEArec are complementary tools designed with the same end goal, i.e., improving spike sorting results, but both targeted at a different primary audience.

## Information Sharing Statement

The presented graphical user interface and spike sorting analysis code are available under an open-source license from https://github.com/jwouters91/shybrid. The software is also available as a Python package from https://pypi.org/project/shybrid/. The extracellular traces (2015 09 03 Cell.9.0) that were hybridized and analysed in this work were collected by [24] and are available from their website http://www.kampff-lab.org/validating-electrodes.

## Acknowledgment

The authors would like to thank Jonathan Dan and Jonathan Moeyersons for their time spent on thoroughly testing the software and for their valuable feedback.

## Appendix

## A. Auto hybridization fitting factor bounds

The calculation of the fitting factor bounds during the automatic hybridization is based on robust statistics, which are commonly used for the detection and removal of outliers. The automatic bounds selection is rather conservative, i.e., it is likely that quite a few good spikes are excluded from the hybridization when using the automated approach.

Consider 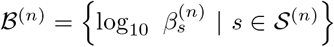 which is the set of the logarithm of the fitting factors (see Section II) for a certain neuron *n*. The logarithm is used to be able to also remove close to zero fitting factors based on simple statistics. Given 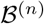, the first and third quartile are calculated, denoted by Q_1_ and Q_3_ respectively. From those quartile values the interquartile range (IQR) is calculated as IQR = Q_3_ - Q_1_. From those statistics the bounds are calculated:

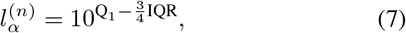

and

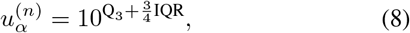

where the IQR scaling factor (i.e. 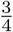) was determined experimentally.

## B. Auto hybridization random unit relocation

During the automatic hybridization, a random unit relocation is calculated for every neuron. For this relocation, only a shift in the y-direction is considered. The random shift is determined by drawing a y-position on the probe grid model (see Section II) from a discrete uniform distribution. This random y-position is the y-position to which the channel with the maximal deflection in the spike template is shifted to. In this way we avoid that the complete template is shifted off the probe. The actual shift can then be calculated as the random y-position minus the y-position of the channel with maximal deflection in the original template. A minimum shift of two channels is enforced, to make sure that the re-inserted unit is sufficiently separable from the original unit.

## C. External template import

When an external template is imported, there are no spike times available, neither is the scaling known. The spike occurrences are modeled as a poisson point process. The inter-spike interval Δ_ISI_ is then modelled by drawing from an exponential distribution:

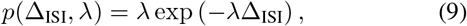

where λ represents the desired spike rate. Every inter-spike interval sample 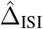 is enforced to last at minimum the user-defined refractory period Δ_min_:

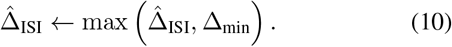

The actual simulated discrete spike times ksim are obtained by calculating the cumulative sum over the inter-spike interval samples. Those spike times are then discretized by multiplying them with the recording sampling frequency and rounding each product to its nearest integer. This gives rise to a set of discrete spike times 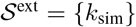.

The template scaling is derived from the user-defined desired peak-signal-to-noise ratio 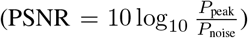. The scaling factor is calculated as follows:

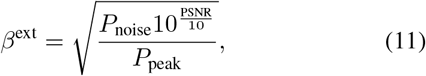

with *P*_peak_ equal to the square of the peak absolute value over all channels of the external template and *P*_noise_ equal to a robust estimate (based on the median absolute deviation) of the noise variance of the channel on which the template reaches its peak absolute value.

The hybrid data generated from an external template can then be described as follows:

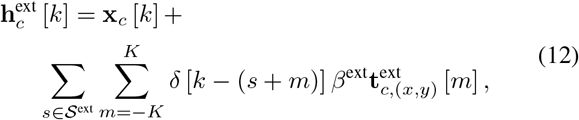

where 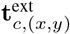 denotes the imported external template at channel *c*. Note that the template temporal window is derived from the external template directly. The external template is assumed to match the sampling frequency of the recording data that is being hybridized.

## D. Automatic merging

The merging framework for a specific ground-truth spike train consists of the following steps:

1. Compute the correspondence between the ground-truth spike train and all automatically recovered spike clusters in terms of precision and recall. More information on those performance metrics can be found in Section III-G2.
2. Sort all clusters on descending precision, such that the cluster with the highest fraction of true spike times is on top of the list.
3. Merge the ordered clusters together in a top-down fashion, i.e. starting from the cluster with the highest precision, as long as the merge operation increases the F_1_-score of the new cluster that contains all previously merged clusters.

Initially, the merging of clusters with a high precision will increase the sensitivity, at only a very small drop in precision. Such a merging will likely lead to an increase in F_1_ -score. At a certain point, clusters will start containing significant amounts of false positives that will notably decrease the precision of the merged cluster. This decrease will then result in a decreasing F_1_-score. The proposed approach tries to find the combination of clusters with maximal F_1_ -score, without explicitly having to consider all possible combinations, preventing a combinatorial explosion from happening.

## Footnotes

1 The tool is available on https://github.com/jwouters91/shybrid.

2 Please consult the phy documentation for more information about the template-gui format.

